# Neuronal morphology and projection analysis by multispectral tracing in densely labeled mouse brain

**DOI:** 10.1101/230458

**Authors:** Douglas H. Roossien, John M. Webb, Benjamin V. Sadis, Yan Yan, Lia Y. Min, Aslan S. Dizaji, Luke J. Bogart, Cristina Mazuski, Robert S. Huth, Johanna S. Stecher, Sriakhila Akula, Jeff W. Lichtman, Takao K. Hensch, Erik D. Herzog, Dawen Cai

## Abstract

Accurate and complete neuronal wiring diagrams are necessary for understanding brain function at many scales from long-range interregional projections to microcircuits. Traditionally, light microscopy-based anatomical reconstructions use monochromatic labeling and therefore necessitate sparse labeling to eliminate tracing ambiguity between intermingled neurons. Consequently, our knowledge of neuronal morphology has largely been based on averaged estimations across many samples. Recently developed second-generation Brainbow tools promise to circumvent this limitation by revealing fine anatomical details of many unambiguously identifiable neurons in densely labeled samples. Yet, a means to quantify and analyze the information is currently lacking. Therefore, we developed nTracer, an ImageJ plugin capable of rapidly and accurately reconstructing whole-cell morphology of large neuronal populations in densely labeled brains.

## Introduction

Despite neural circuits being of wide interest to neuroscientists, there are few technologies that fully dissect these complex networks. Automated serial electron microscopy techniques are one means of obtaining the full details of networks, but the volumes that can be studied are limited by the throughput or other physical constrains of the techniques being employed, therefore currently limited to smaller vertebrates^1^. While light microscopy is easy to implement and allows for imaging large volumes of genetically labeled neurons, the inability to distinguish processes between labeled neurons is a considerable limitation, such that reliable anatomical neuronal reconstruction results can only be obtained from samples in which labeled neurons occupy unique volumes with little to no overlap, or, alternatively, singly labeled neurons^2–7^. Generating such samples can be cumbersome and increases the number of samples needed to obtain statistically relevant results. One approach to overcome the limitation of light microscopy is to differentially label neighboring neurons with distinct colors. Brainbow does so by expressing random ratios of three or more fluorescent proteins (FPs) through Cre/Lox recombination in specific populations of neurons within a single brain ^8,9^. These unique colors make identifying multiple neurons in a densely-labeled sample possible. In addition, because FPs from second generation Brainbow reagents are membrane bound and compatible with immunofluorescence amplification they allow visualization of fine morphological details such as dendritic spines and axonal boutons. In turn this permits identification of putative synaptic locations and can uncover anatomical constraints on neuronal circuitry. However, a lack of tools for quantitative analysis has limited Brainbow application.

Currently, popular commercial software such as Neurolucida and iMaris allow either manual tracing or user-guided tracing algorithms, but none handle multispectral images^6^. We thus wrote nTracer, a java-based program to facilitate post-acquisition processing and user-guided semi-automated tracing of Brainbow multispectral images. We then trace a variety of different neuronal subtypes including cholinergic neurons in the striatum, granule cells in the dentate gyrus, vasoactive intestinal polypeptide (VIP) expressing neurons in the suprachiasmatic nucleus and parvalbumin (PV) expressing basket cells in the hippocampus and visual cortex. We found tracing results with nTracer can be generated at an average rate of a few hours per neuron. Together this suggests nTracer produces rapid and accurate tracing of densely labeled Brainbow samples. Overall, nTracer is an important tool that will allow neuroscience researchers to analyze morphology and anatomy of large populations of neurons within single samples using the light microscope.

## Methods

### Stereotaxic injection

The two Brainbow AAV (AAV9.hEF1a.lox.TagBFP.lox.eYFP.lox.WPRE.hGH-InvBYF and AAV9.hEF1a.lox.mCherry.lox.mTFP1.lox.WPRE.hGH-InvCheTF) were obtained from the UPenn Vector Core and mixed to equal titer (8 x10^12^). All surgeries were performed in accordance with the institutional animal guidelines and approvals of the University of Michigan, Harvard University, University of Washington at St. Louis and the Massachusetts Institute of Technology. Mice were anesthetized with either a mixture of ketamine (90mg/kg) and xylazine (9mg/kg) or Avertin (250mg/kg) or continuously delivered isoflurane and mounted on a stereotaxic frame (WPI #502600). Craniotomies were performed using a small dental drill and injections were performed using a 30 gauge stainless steel needle inserted directly through the dura^10^.

Final Brainbow virus volume (µL) x titers (in GC/mL) and stereotaxic coordinates (in mm from Bregma/dura AP, ML, DV) used in this study are as follows: ChAT-Cre (Chat^tm2(cre)Lowl^ Jackson Laboratory No 006410; Rossi et al 2011 – Cell Metab 13(2):195-204) 1µL x 8×10^11^, (+1.1, −2.0, −3.4); POMC-Cre (hypothalamus) (Tg(Pomc1-cre)16Lowl Jackson Laboratory No 005965; Balthasar et al 2004 – Neuron 42(6):983-91) 1µL x 1×10^12^, (−2.2, −0.25, −5.4); POMC-Cre (dentate gyrus) 1µL x 1.6×10^12^, (−2.0, −1.6, −2.0); PV-Cre (V1) (Pvalb^tm1(cre)Arbr^ Jackson Laboratory No 008069; Hippenmeyer et al 2005 – PLoS Biol 3(5):e159) 1uL x 3×10^12^ (−3.0, ±2.0, −0.4), PV-Cre (CA1) 1µL x 8×10^11^, (−2.0, −1.8, −1.6); VIP-Cre (SCN) (Vip^tm1(cre)Zjh/J^ Jackson Laboratory No 010908; Taniguchi et al 2011 Neuron 71:995-1013) 1µL x 1×10^12^ or 2.5×10^11^, (+0.4, −0.2, −5.5); Wildtype CD1 (CA2) AAV.Brainbow 1µL x 1.6×10^12^ + AAV.CaMKII-Cre 1×10^11^ (−1.1, −1.0, −1.9). At least 3-4 weeks after injection, mice were perfused under heavy anesthesia with cold PBS followed by cold 4% paraformaldehyde (PFA). Brains are dissected immediately following perfusion and post-fixed in 4% PFA at 4°C with gentle shaking for 24 hours.

### Immunohistology

We chose non-cross-reactive primary antibodies raised from 4 distinct animal species to specifically recognize the 4 FPs expressed by Brainbow AAV^9^: rat anti-mTFP1, chicken anti-EGFP, rabbit anti-mCherry, and Guinea Pig anti-mKate2 (available from Kerafast, Boston MA). The mouse anti-GAD65 primary was obtained from the Developmental Studies Hybridoma Bank (Gad-6) and used at 1:500 dilution. The mouse anti-PV antibody was obtained from Swant (#235) and used at a 1:500 dilution. Four non-cross-reactive secondary antibodies, conjugated with spectrally well-separated fluorophores, were chosen for the Brainbow primaries to allow optimal recording of enhanced fluorescence in each spectral channel throughout the brain sections: anti-Rat Alexa594 (Life Technologies #A21209), anti-Chicken Alexa488 Jackson ImmunoResearch Inc #703-545-155), anti-Rabbit Alexa546 (Life Technologies #A10040), and anti-Guinea Pig Alexa647 (Jackson ImmunoResearch Inc #706-605-148). The GAD65 and PV stains were visualized with anti-mouse 405.

Sections were first blocked and permeabilized in StartingBlock-PBS (ThermoFisher) with 0.5% Triton X-100 for 2-4 h at room temperature (RT) with gentle shaking. After wash, sections were incubated in all four primary antibodies each with a dilution of 1:500 in PBS with 0.25% Triton X-100 (PBST) and 0.2% sodium azide for 7-10 days at 4°C with gentle shaking. Primary antibodies were washed out of sections with three changes of 0.25% PBST for 1 hr each at RT. Secondary antibodies were mixed and diluted to a final concentration of ~3-4 ug/mL in 0.25% PBST with 0.2% sodium azide and added to sections for 3-5 days at 4°C with gentle shaking. After secondary incubation, two washes in 0.25% PBST for 2 h each followed by a final 2 h wash in PBS were performed. Sections were either mounted in Vectashield (Vector Labs #H-1000) or treated with a gradient of fructose solution up to a final concentration of 70% (w/w) for refractive index matching^11^.

### Imaging

Sections were imaged on a Zeiss LSM 780. Optimal separation of fluorophores was achieved using a 488nm laser for Alexa488, a 543nm laser for Alexa546, and a 633nm laser for both Alexa594 and Alexa647 together with a fixed 488/543/647 dichroic mirror. Alexa488, Alexa594 and Alexa647 were imaged simultaneously and fluorescence was collected in 3 separate channels. Alexa546 was imaged in a subsequent scan with fluorescence collected in a 4^th^ channel. Objectives were either 20x W Plan APOCHROMAT NA 1.0 water or 40x C Plan APOCHROMAT NA 1.3 oil objective. While these settings are optimized for spectral separation, due consideration has also been given to scan time. For example, better spectral separation could be obtained by imaging Alexa594 with a 594 laser on a separate track. However, this would significantly increase scan time due not only to the need for an additional independent scan, but also for the need to switch the dichroic between twice for each frame. The setting we used here was much faster, but results in significant Alexa594 emission being collected in channel 3 as well as channel 2. Therefore, linear unmixing was performed to separate these two fluorescent emissions^12^. The laser intensity correction in depth function was used in the Zen software was used to compensate for light scattering and photobleaching.

## Results

### Development of post-acquisition processing functions

Reliable tracing of Brainbow labeled neurons depends on obtaining high contrast and chromatic error-free 3D images. In other words, image stacks need to contain neurons labeled in distinct and consistent color in three dimensions. Here we define color (or color signature) as the composite of intensity values from each channel for every given pixel. For example, a specific neuron in 8-bit may have a color signature of Ch1:156, Ch2:005, Ch3:255, Ch4:073. In biological terms this amounts to the ratio of FP expression in each neuron. Due to the scattering nature of the brain and the inherent optical aberrations of confocal microscopy, however, Brainbow images often decrease in quality and contain color defects when imaging deep into the tissue. We therefore first developed post-acquisition processing functions in nTracer that correct for these color defects and facilitate analysis of neuronal networks through >100μm thick tissue sections. These have been assembled into one convenient ImageJ User Interface called AlignMaster included in the nTracer software package. Figure 1 illustrates an optimized general workflow for Brainbow labeling with second-generation Brainbow reagents^9^, imaging, post-acquisition processing and neuronal tracing (**Fig. 1**).

**Figure 1.**
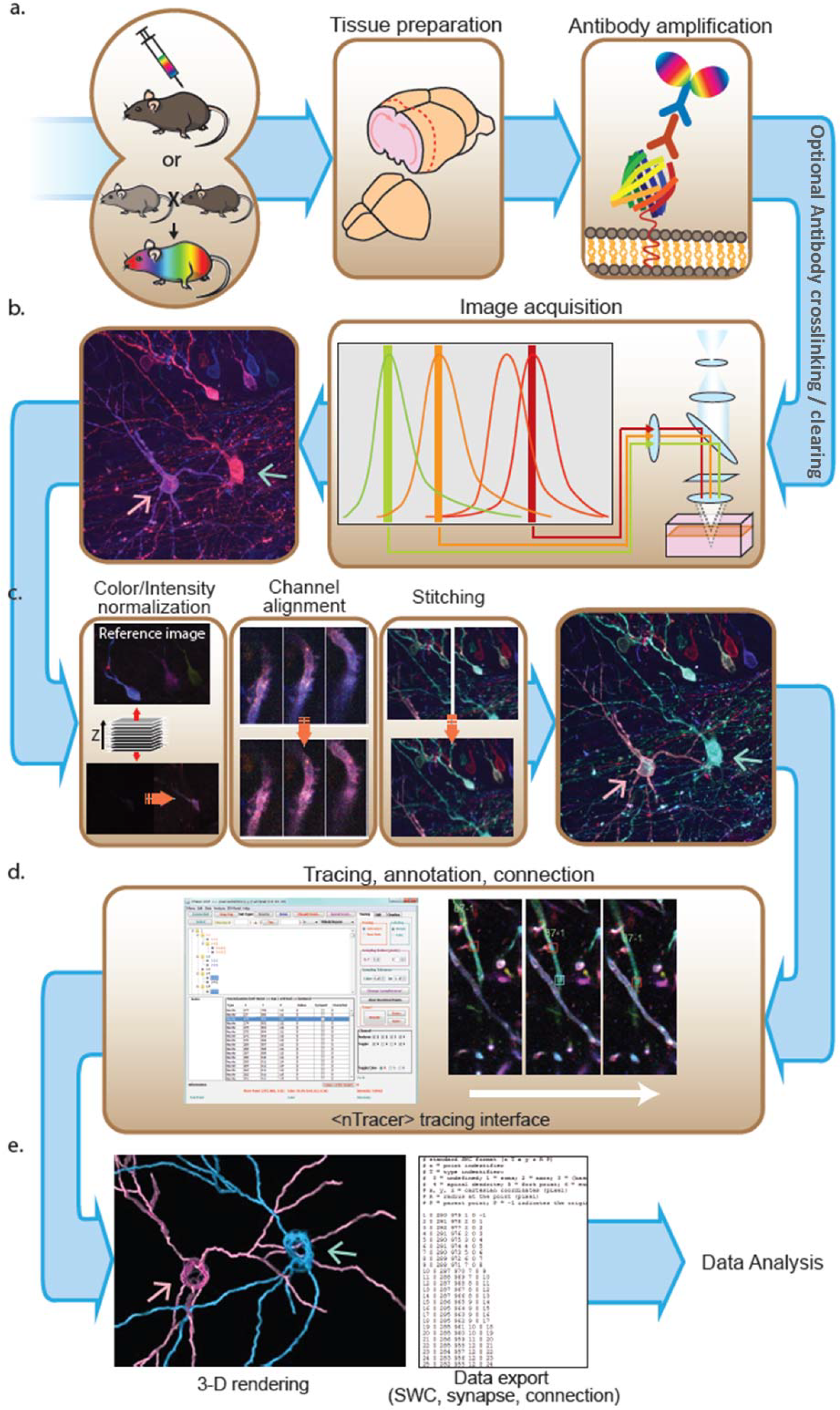
Flow chart for Brainbow labeling, imaging and tracing for neural circuit reconstruction. (a) To accurately reconstruct the projections from many neuronal somata to their terminals, we used second generation Brainbow reagents, including adeno-associated viruses (Brainbow AAVs) and transgenic mouse lines (Brainbow3.x) because their farnesylated fluorescent proteins (FPs) label homogenously throughout the whole neuron including the finest dendritic spines and neuronal processes. (b) Four non-cross-reactive secondary antibodies conjugated with spectrally well-separated fluorophores allow optimal recording of enhanced fluorescence in each spectral channel throughout the brain sections. (c) To correct imaging defects in post-acquisition processing, nTracer package provides functions to automatically 1) normalize image intensity across all spectral channels and z-depths; 2) perform channel registration to correct optical misalignment and chromatic aberration of the microscopy system and 3) rapidly stitch image tiles to create a final image in which each neurons are labeled by rich and consistent colors throughout the image stack. (d) nTracer as an ImageJ plugin package allows user-guided tracing of neurons in distinct colors in densely labeled samples. (e) The tracing results are exportable in various formats for 3D line art rendering, single cell morphology and connectivity analysis.

There are two major sources of color defects when imaging Brainbow samples. The first is due to absorption, scattering and photobleaching, which causes a gradual decrease in fluorescence intensity when imaging deeper into the tissue. Although gradually increasing excitation power and/or detection gain can partially compensate this intensity drop, it becomes practically impossible to maintain constant intensity throughout the image stack. Compounding this is the fact that the intensities of each FP will decrease at different rates/amounts, resulting in inconsistencies in color signature at different z-positions (**Suppl. Fig. 1a**). To address these concerns, we added a histogram matching correction to nTracer to normalize signal intensity while maintaining the intensity ratios between channels. The user is asked to select a reference image slice to which nTracer matches the histogram (see below) of each slice in the image stack. While the reference slice can be chosen from any channel or focal plane, the optimal reference will contain an evenly distributed histogram with minimal pixel values greater than 95% of the maximum bin (**Suppl. Fig. 1c**). This ensures that FP intensity remains constant in any depth of the 3D stack while minimizing amplification of background noise (**Suppl. Fig. 1c**). The reference image’s cumulative probability distribution function *CDF_ref_*() of its histogram *H_ref_*() is calculated. For each target image slice in all channels of the whole 3D stack, a new histogram *H_tar_*() is applied, which satisfies the condition that for each gray scale level *G_ref_* a *G_tar_* is determined to satisfy *CDF_ref_*(*G_ref_*) = *CDF_tar_*(*G_tar_*). After intensity equalization, we found the histogram of all image slices to be similar to that of the reference image.

The second source of color defects results from imperfect optical alignments and chromatic aberrations in the microscope system. Such defects cause spatial misalignment between spectral channels that creates inconsistent color along the edge of many neuronal processes (**Suppl. Fig. 1b**). We also corrected for this in the nTracer package, by designing a channel registration function. In brief, the user makes a point selection on the image where a neuronal feature appears in all spectral channels; and then nTracer determines the translational shifts that correspond to the greatest masked-intensity correlations between each channel and the selected channel. Eliminating highly correlated background pixels, the masking function increase the sensitivity of correlating the fluorescent neurons. This correction is particularly useful for microscope systems with large chromatic aberration, whose severe par-focal problem normally causes large shifts between short and long wavelength channels (**Suppl. Fig. 1d**).

In addition to these corrections, nTracer provides a 3D stitching function that allows rapid merging from overlapping Brainbow image tiles to create a single image stack that covers a large tissue volume. While the size of datasets can vary dramatically depending on the biological application, the size of the datasets used throughout this report varies from hundreds of megabytes to hundreds of gigabytes.

Both the intensity correction and channel alignment can be done in batch mode on all images (e.g. all stacks if a multi-tile image was taken for stitching) in a selected folder. For stitching, easily identifiable features in overlapping regions are selected by the user and local area is used to calculate the translation parameters needed to stitch the tiles together. To minimize processing time, which can be considerable with larger data sets, both the alignment and stitching functions rely on a local area to perform initial translation calculations and previews to proof the result prior to applying the translation. The data sets in this study were first processed with the intensity correction, followed by channel alignment, then stitching. We found it important to do batch processing of color intensity prior to stitching to obtain uniform signal-to-noise and intensity across all tiles. Additional downstream processing steps can be performed as needed. For example, deconvolution using the Richardson-Lucy algorithm in DeconvolutionLab^13^ was performed on the images shown in Fig. 2c.

**Figure 2.**
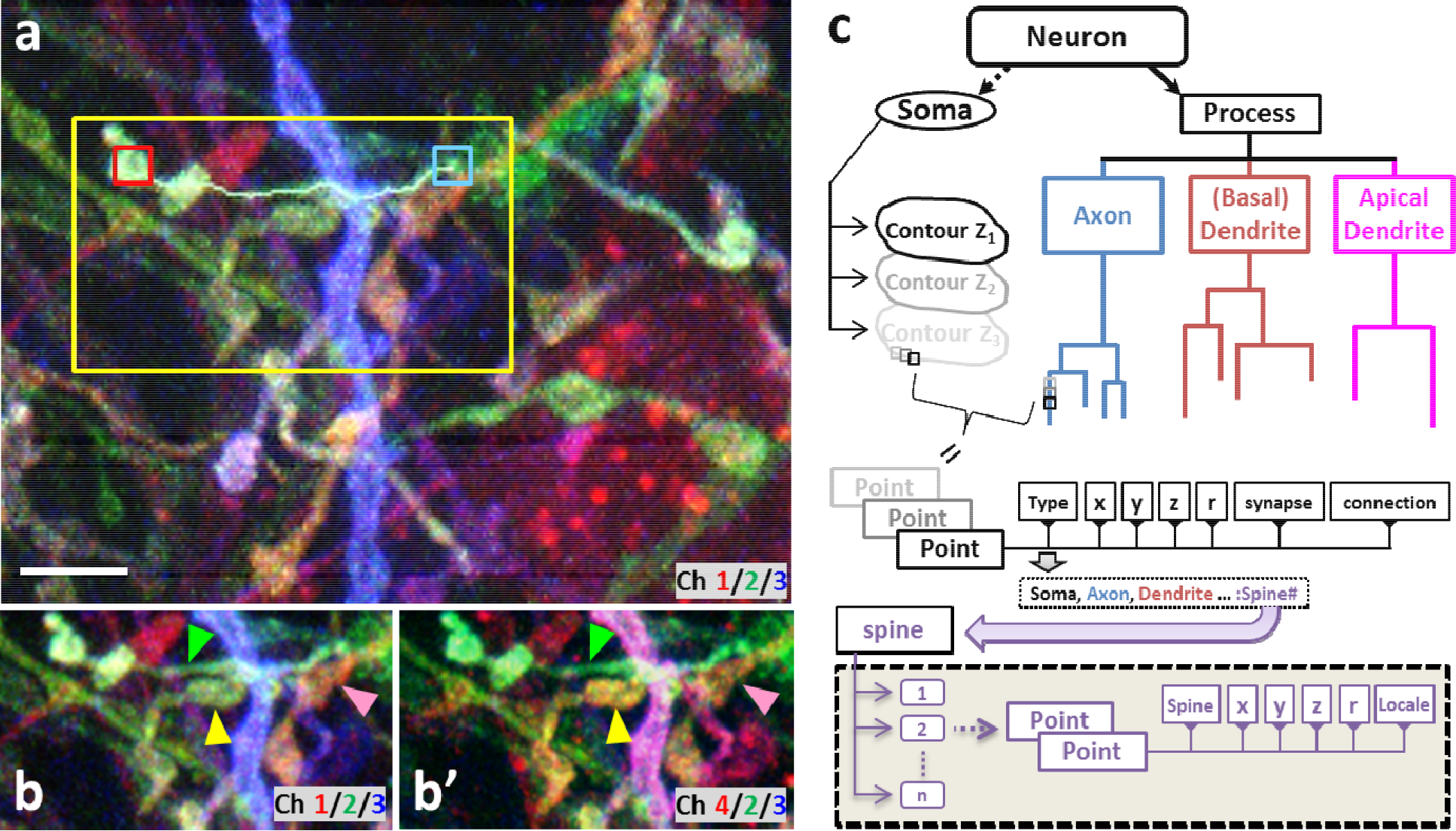
nTracer basic functionality and data structure. (a) To start tracing, a mouse-click is placed in the vicinity of the neurite to be traced. nTracer utilizes the color profile and a “mean-shift” algorithm to accurately reposition this mouse-input onto the neurite as the start point (red box). The tracing end point (cyan box) is determined in the same way with additional constrains to make sure that the end point has a similar color profile. A keyboard hotkey is then used to trace as a neurite/spine in 3D. (b-b’) Two sets of RGB composite images (best color separation for human tri-chromatic vision) can be toggled in nTracer to display 4 spectral channels taken from the 4-FP Brainbow AAV labeled samples. As not all the spectral information is displayed at the same time, it is possible that neurites appearing the same color actually belong to different neurons (compare green to yellow and yellow to pink arrowheads in b and b’). Regardless of how the images are displayed, nTracer uses information from all four channels to prevent tracing errors arising from human vision limitations. (c) Diagram of nTracer results data structure as detailed in Methods. Scale bar is 5 μm.

### Tracing function and data structure

The current bottleneck in fully automated tracing algorithms is proof reading and error correcting (refs 6-8). Because of this, combined with the lack of any ground-truth Brainbow tracing results for validation, we developed nTracer as a user-guided semi-automated tracing software which allows for “on-the-fly” editing of tracing results. Our approach was to design a system where user-defined anchor points on a neurite are joined by a rapidly generated skeletal tracing. To make this tool widely available and easy to use we configured it as an ImageJ plugin^14^.

We incorporated two algorithms into nTracer to accurately trace neurites from specific neurons. The first prevents user error by not allowing anchor points to be assigned to different neurons, and the second prevents the skeletal trace from jumping to the wrong neuron when joining anchor points. To start a tracing, the user identifies a neuron to be traced and uses a mouse click to suggest a start point on its process and to measure the neuron’s color signature around the start point. In most cases, the mouse clicks hardly land onto the “right” spot, which results in inaccurate color sampling. nTracer solves this problem by applying a mean-shift algorithm to automatically refine the user input and settle the start point onto the center or membrane wall of the targeted neurite with high labeling intensity^15^ (**Suppl. Fig. 2**). The end point is defined in a similar way with additional constraints set by the color signature sampled around the start point. The user can therefore avoid setting an end point onto a different neuronal process due to human visual or computer display limitations, in particular with Brainbow images composing of more than 3 spectral channels (see main text and **Suppl. Fig. 3**). To generate a smooth track along the neuronal process, nTracer utilizes the A* algorithm^16^ to connect the two anchor points with a least-cost path, similar to that implemented in “Simple Neurite Tracer”, which was designed for tracing monochromic images^17^. nTracer defines the A* cost at voxel *i* as a weighted sum of the normalized spectral and intensity difference between the start point *p* and voxel point *i*, which can be formulated as:

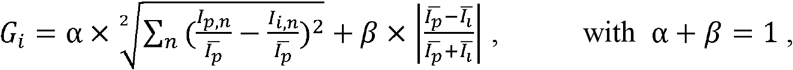

where *n* denotes the n^tn^ spectral channel. 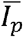 and 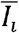 are the total intensities in all spectral channels at start point *p* and voxel point *i*, respectively. By constraining the pathfinding range to the voxels enclosing the two anchor points and by choosing optimized heuristic values calculated based on windowing-smoothed voxels, nTracer creates an optimal minimal path almost instantaneously, while variance thresholds ensure that any path containing large intensity or color gaps will be rejected. In addition, the tracing process with nTracer is iterative, reducing user burden by only requiring one click per trace after the initial trace on a neurite.

**Figure 3.**
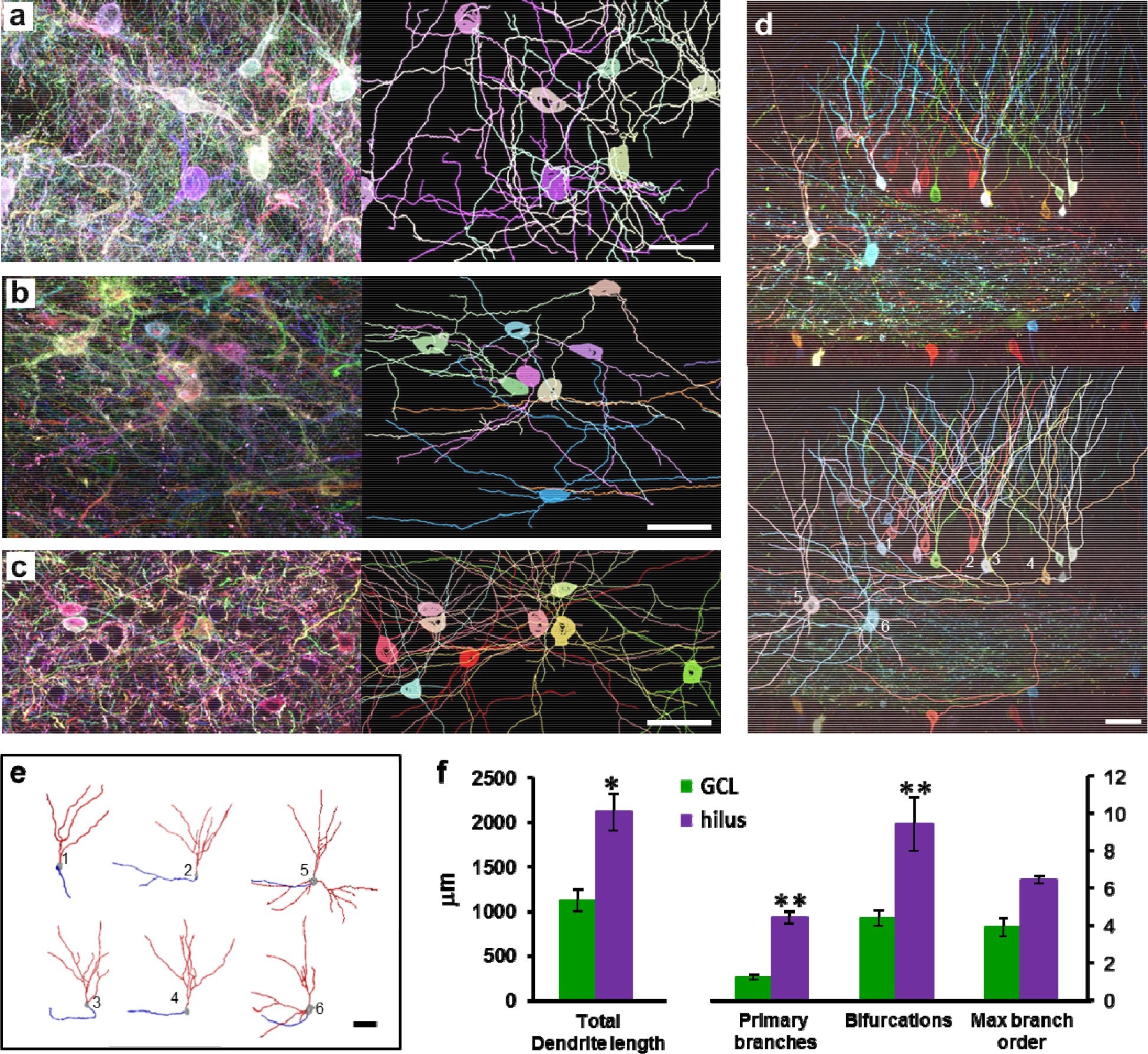
Morphological analysis of single cells in densely labeled samples by nTracer. (a-d) Paired maximum z-projections from AAV.Brainbow-injected data sets and subsequent nTracer results shows the versatility of nTracer for studying diverse cell types in multiple brain regions. Representative reconstructions of (a) cholinergic cells from ChAT-Cre mouse injected in dorsolateral striatum, (b) POMC neurons from POMC-Cre mouse injected in arcruate nucleus of the hypothalamus, (c) basket cells from PV-Cre mouse injected in visual cortex, and (d) granule cells from POMC-Cre mouse injected in dentate gyrus. (e) Individual granule cells from (d) were exported from tracing results and displayed with color-coded compartments (red = dendrites, blue = axons). Here, cells 1-4 were localized in the granule cell layer (GCL) while cells 5-6 were localized in the hilus. (f) Granule cells localized in the hilus had longer total dendrite length, more primary branches and bifurcations than those localized in GCL; but they all had the same maximum branch order. Note that all granule cells used for analysis were taken from the same dentate gyrus section. All values displayed as mean ± SEM. * p < 0.05, ** p < 0.01 by unpaired two-tailed Student’s t-test. Detailed single-cell morphometric analyses of all of the cells in this figure can be found in Table 1. Scale bars are 50 μm.

**Table 1.**
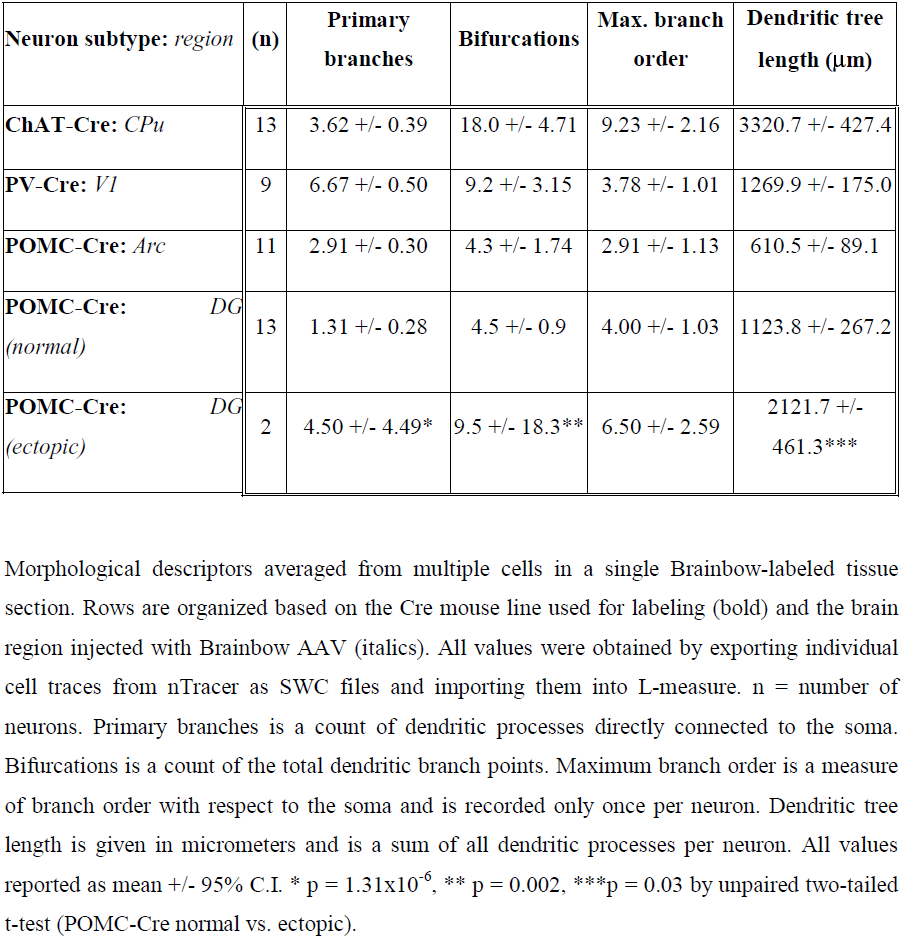
Single cell morphometric analysis of dendritic trees.

Because nTracer must record multiple neurons to reconstruct complex circuits in the same image, it uses a data structure that differs from software approaches that focus on tracing single neurons from one image at a time. nTracer utilizes the generic JTree structure of JAVA to allow flexible storage and modification of tracing points of multiple neurons in the computer memory. Three JTrees are built for each traced cell to store the tracing points of the somas, processes and spines independently (**Fig. 1d** and **Fig. 2c**). The soma tree contains parallel nodes, each of which stores soma tracing points on a Z plane. The process tree contains parallel nodes, each of which is a bifurcated branching tree that stores connected branches of an axon, or a dendrite. The soma contour or neurite branch is composed of connected tracing points, each of which is a 7-element data array containing the type of the tracing point (Soma, Dendrite, Axon, Spine etc.), x, y, z coordinates, radius at the point (0 for a soma point or for where the process radius is not determined), whether or not a synapse, and its connection status. Spines can also be traced off from a dendrite or soma point (has a type of Spine) and stored as parallel non-branching nodes in the third tree-structure database. Each spine tracing point is a 6-element data array that stores the type (Spine), x, y, z coordinates, radius at the point, and its locale information (soma or dendrite name). The tracing results (including connectivity information), raw image information and nTracer setting parameters can be saved in files of custom format and exported as line art image stacks for volume rendering (**Suppl. Fig. 4 and Video 5-10**). Tracing results of each neuron can also be exported as separate files in standard SWC format^18^ for morphology analysis and rendering with other software, such as L-measure^19^.

**Figure 4.**
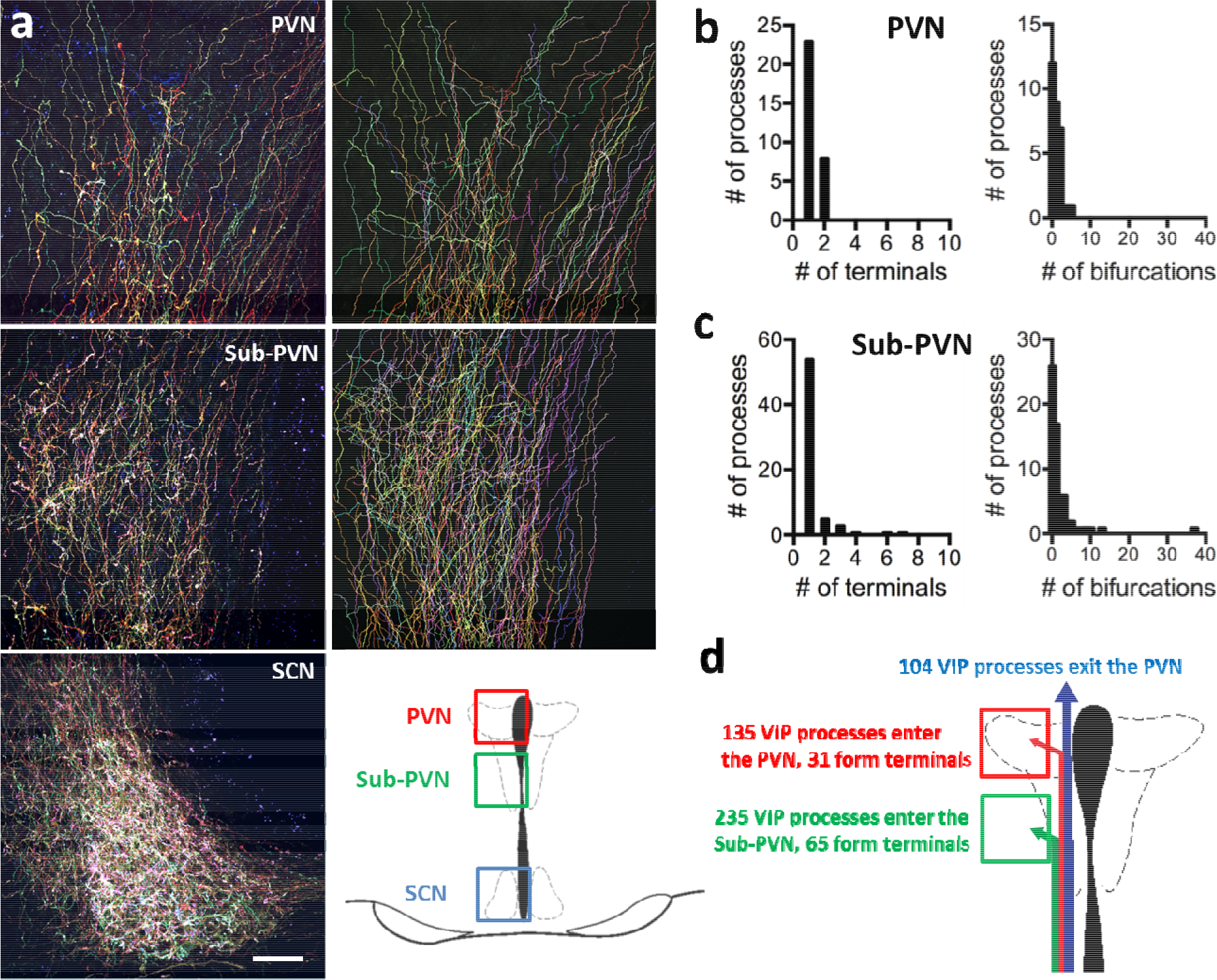
Projection analysis of SCN VIP divergence in the sub-PVN and PVN. (a) Unilateral labeling of SCN VIP neurons using Brainbow AAVs reveals processes in the sub-PVN and PVN (left panels). Using nTracer (right panels), more than 200 processes can be mapped from a single neuronal population in one brain section. Anatomical characteristics of these processes can be quantified: (b and c) show the number of terminal synapses (left panels) and bifurcations per process (right panels) of VIP processes entering PVN and sub-PVN, respectively. (d) A schematic details the number of VIP processes identified in the sub-PVN and PVN and the fraction that form terminals. Scale bar is 50 μm.

### Mapping projection patterns of a complete population of neurons from a single Brainbow sample

We next sought to demonstrate the utility of nTracer. We chose to focus on the vasoactive intestinal polypeptide-expressing (VIP) neurons in the hypothalamic suprachiasmatic nucleus (SCN). These neurons project dorsally to form synaptic terminals in the paraventricular and sub-paraventricular nuclei (PVN and sub-PVN, respectively^20^), although it is not known whether individual VIP neurons project to multiple PVN targets or send convergent inputs onto single PVN targets. We injected Brainbow AAVs into the SCN of one hemisphere of mice expressing Cre under control of the VIP promoter (VIP-Cre^21^). We found that injection of virus at 1:5 (~1×10^12^ GC/ml; n=2 brains) to 1:20 dilution (~2.5×10^11^ GC/ml; n=2 brains) resulted in labeling of similar numbers of axons projecting dorsally with 1:20 producing discriminable colors and 1:5 producing saturated labeling. We therefore present results from the brains injected with the more diluted virus (**Fig. 4**). In both brains, we found many axons and a small number of cell bodies labeled in the SCN contralateral to the side of injection. We used nTracer to trace all of the individual dorsal projections in the bilateral sub-PVN and PVN, an area of more than 1.3 mm x 0.6 mm, and performed further analysis of neurites with at least one terminal in the PVN or sub-PVN (**Fig. 4b-c**). We found that the large majority of neurites (~75%) bifurcate only 0 to 1 times while only a few VIP neurons had processes with up to 10 branch points outside the SCN. These results indicate that individual SCN VIP neurons each target a small population of cells in the PVN or sub-PVN.

## Discussion

Included in the nTracer package are image processing tools designed to optimize tracing including correction of intensity drifting in depth due to uneven illumination, scattering, and photobleaching; correction of channel misalignment due to hardware misalignments and optical aberrations in the microscopy system; and intensity normalization and merging of overlapping adjacent image tiles. The key advantage of nTracer is its ability to trace neurons based on color signature, making it compatible with densely labeled Brainbow samples and to fully actualize this powerful neuronal labeling technique.

For this study we chose mouse Cre lines and injection sites in order to label regions with a variety of gross anatomical features. For example, the hippocampus and cortex have distinct laminar cellular organization whereas the hypothalamus and striatum do not. Labeling an entire Cre+ population of neurons in a region of interest is attainable with a wide titer range of Brainbow AAVs (from ~1×10^10^ to 1×10^13^ GC/ml), however higher (~10^13^ GC/ml) or lower titers (~5×10^10^ GC/ml) resulted in more neurons labeled in saturated colors (white cells) or simple colors (expressing only one species of FP), respectively. This is directly dependent on the density of the neuronal subtype in the region of interest and the strength and fidelity of the Cre driver. Therefore, a limitation to our overall approach is the need to determine injection coordinates and viral titers empirically for each sample type. However, because we rely on immunolabeling of samples our approach is compatible with other standard histological techniques. For example, cell-type specific markers can be used in combination with Brainbow staining to trace multiple cell types within a circuit or, alternatively, axonal or dendritic markers can be used to distinguish neurite type in neurons with unknown or less stereotypic morphologies.

Because nTracer is user-guided and not fully automated, it requires significant man-hours to trace complete data sets. While semi-automated tracing algorithms used in other software, such as Neurolucida, do not require as much user attention to generate the tracing result, there remains a bottleneck in the need for proofing the results due to the propensity for tracing errors^22,23^. Since the error rate increases substantially with more neurons and greater density, eventually the effort spent on proofing and post-tracing editing outweighs the benefit gained from semi-automated tracing. With nTracer, the tracing results are displayed immediately and therefore proofing and editing is done “on-the-fly”.

In the samples that all VIP-neurons were labeled, we were not able to map connectivity within the SCN or to follow the projections back to the cell body due to color blending between neuronal processes that are close to each other. This is due to the optical resolution limitations of confocal microscopy. Combining Brainbow labeling with emerging super-resolution light microscopy techniques, such as Expansion Microscopy^24^ and its variant protein-retention ExM^25,26^, is an exciting possibility for producing images at the spatial resolutions that are suitable to distinguish the closely positioned synapses and neuronal processes. Going forward, we expect the combination of Expansion Brainbow Microscopy and nTracer will allow full tracing of complete neurons in SCN.

Nonetheless, we are currently working on developing additional semi-automated and automated tracing algorithms for nTracer^27^. The tracing results developed with the current nTracer described here will serve as a ground-truth for their development. nTracer will also benefit from adapting the more versatile data structure of the ImageJ2 platform currently under development^28^.

## Software

Fiji/ImageJ plugins and manual for nTracer and AlignMaster (post-imaging processing) are available at https://www.cai-lab.org/ntracer-tutorial.

## Acknowledgements

We would like to acknowledge Carl Zeiss Microscopy for the LSM780 confocal microscope, H. Akil and W.T. Dauer for the PomC-Cre and ChAT-Cre mice, respectively. This work was supported by Michigan miBRAIN initiative to Y.Y. and D.C., by NIH/NIAI (R01AI130303) and NSF/Neuronex-MINT (NSF-1707316) to D.H.R and D.C., and by NIH/NIMH (R01MH110932) to D.C. This work was supported by NIH (F31GM116517) to C.M. and NIH/NINDS (R01NS095367) to E.D.H. T.K.H and J.W.L were supported by NIH/NIMH Silvio Conte Center (P50MH094271). J.W.L. was supported by NIH/NINDS (DP2OD006514, R01NS076467, U01NS090449 and P41GM10371), MURI Army Research Office (W911NF1210594 and IIS-1447786).

## References

1. Hildebrand, D. G. C. et al. Whole-brain serial-section electron microscopy in larval zebrafish. Nature 545, 345–349 (2017).

2. Chothani, P., Mehta, V. & Stepanyants, A. Automated tracing of neurites from light microscopy stacks of images. Neuroinformatics 9, 263–78 (2011).

3. Osten, P. & Margrie, T. W. Mapping brain circuitry with a light microscope. Nature Methods 10, 515–523 (2013).

4. Meijering, E. Neuron tracing in perspective. Cytom. Part A 77, 693–704 (2010).

5. Peng, H. et al. BigNeuron: Large-Scale 3D Neuron Reconstruction from Optical Microscopy Images. Neuron 87, 252–256 (2015).

6. Parekh, R. & Ascoli, G. A. Neuronal morphology goes digital: a research hub for cellular and system neuroscience. Neuron 77, 1017–38 (2013).

7. Lu, J. Neuronal tracing for connectomic studies. Neuroinformatics 9, 159–166 (2011).

8. Livet, J. et al. Transgenic strategies for combinatorial expression of fluorescent proteins in the nervous system. Nature 450, 56–62 (2007).

9. Cai, D., Cohen, K. B., Luo, T., Lichtman, J. W. & Sanes, J. R. Improved tools for the Brainbow toolbox. Nat. Methods 10, 540–7 (2013).

10. Cetin, A., Komai, S., Eliava, M., Seeburg, P. H. & Osten, P. Stereotaxic gene delivery in the rodent brain. Nat. Protoc. 1, 3166–3173 (2006).

11. Ke, M.-T., Fujimoto, S. & Imai, T. SeeDB: a simple and morphology-preserving optical clearing agent for neuronal circuit reconstruction. Nat. Neurosci. 16, 1154–61 (2013).

12. Zimmermann, T. Spectral imaging and linear unmixing in light microscopy. Adv. Biochem. Eng. Biotechnol. 95, 245–65 (2005).

13. Vonesch, C. & Unser, M. A Fast Thresholded Landweber Algorithm for Wavelet-Regularized Multidimensional Deconvolution. IEEE Trans. Image Process. 17, 539–549 (2008).

14. Abramoff, M. D., Magalhães, P. J. & Ram, S. J. Image processing with ImageJ. Biophotonics Int. 11, 36–42 (2004).

15. Yizong Cheng, Y. Mean shift, mode seeking, and clustering. IEEE Trans. Pattern Anal. Mach. Intell. 17, 790–799 (1995).

16. Hart, P., Nilsson, N. & Raphael, B. A Formal Basis for the Heuristic Determination of Minimum Cost Paths. IEEE Trans. Syst. Sci. Cybern. 4, 100–107 (1968).

17. Longair, M. H., Baker, D. a. & Armstrong, J. D. Simple neurite tracer: Open source software for reconstruction, visualization and analysis of neuronal processes. Bioinformatics 27, 2453–4 (2011).

18. Cannon, R. C., Turner, D. a., Pyapali, G. K. & Wheal, H. V. An on-line archive of reconstructed hippocampal neurons. J. Neurosci. Methods 84, 49–54 (1998).

19. Scorcioni, R., Polavaram, S. & Ascoli, G. A. L-Measure: a web-accessible tool for the analysis, comparison and search of digital reconstructions of neuronal morphologies. Nat. Protoc. 3, 866–76 (2008).

20. Teclemariam-Mesbah, R., Kalsbeek, A., Pevet, P. & Buijs, R. M. Direct vasoactive intestinal polypeptide-containing projection from the suprachiasmatic nucleus to spinal projecting hypothalamic paraventricular neurons. Brain Res. 748, 71–6 (1997).

21. Taniguchi, H. et al. A resource of Cre driver lines for genetic targeting of GABAergic neurons in cerebral cortex. Neuron 71, 995–1013 (2011).

22. Liu, Y. The DIADEM and beyond. Neuroinformatics 9, 99–102 (2011).

23. Peng, H., Long, F., Zhao, T. & Myers, E. Proof-editing is the bottleneck of 3D neuron reconstruction: the problem and solutions. Neuroinformatics 9, 103–5 (2011).

24. Chen, F. et al. Optical imaging. Expansion microscopy. Science 347, 543–8 (2015).

25. Chozinski, T. J. et al. Expansion microscopy with conventional antibodies and fluorescent proteins. Nat. Methods 13, 485–8 (2016).

26. Tillberg, P. W. et al. Protein-retention expansion microscopy of cells and tissues labeled using standard fluorescent proteins and antibodies. Nat. Biotechnol. (2016). doi:10.1038/nbt.3625

27. Sümbül, U. et al. Automated scalable segmentation of neurons from multispectral images. in 30th Conference on Neural Information Processing Systems (NIPS) (2016).

28. Schindelin, J., Rueden, C. T., Hiner, M. C. & Eliceiri, K. W. The ImageJ ecosystem: An open platform for biomedical image analysis. Mol. Reprod. Dev. 82, 518–29

